# Proteomic profiling of experimental acute alcohol intoxication reveals FGF21 as a specific ethanol-induced hormone

**DOI:** 10.64898/2026.10.27.762218

**Authors:** Amalie R. Lanng, Samuel A. J. Trammell, Lærke S. Gasbjerg, Matthew P. Gillum, Filip K. Knop

## Abstract

**Background:** The effects of acute alcohol intoxication on the human plasma proteome are still largely unexplored. Gaining insights into this relationship could identify factors that are influenced by and influence alcohol consumption. Fibroblast Growth Factor 21 (FGF21), a hormone produced by the liver and associated with alcohol preferences, is currently undergoing Phase 2 clinical trials for metabolic dysfunction and alcohol-related liver disease. We hypothesized that FGF21 is a key endocrine responsive hormone to alcohol in humans.

**Methods:** In a post hoc analysis, we evaluated the effects of five standard units of alcoholic beverages on the plasma proteome in 12 healthy Caucasian men. Participants, fasting for 10 hours, received 0.7 g of alcohol per kg of body weight via gastric infusion, with blood samples collected before and after consumption.

**Results:** The proteomic response to this level of alcohol intoxication was highly specific. Of 1,462 proteins measured, only 11 changed significantly and FGF21 exhibited by far the largest increase (10.5-fold).

**Conclusion:** This study highlights a unique proteomic response to acute alcohol ingestion and supports further exploration of FGF21 mimetics as potential treatments for alcohol-related disorders.

## Introduction

Alcohol (ethanol) is a natural product of fermentation (e.g., found in fermented fruits and berries) and has been used as a stimulant by humans and their ancestors for millions of years (1). While alcohol is an energy-rich substance (7 kcal/gram vs. 4 kcal/gram for protein and carbohydrate) and, in that context, may promote survival, it also – in high doses – has toxic effects on the human body and results in behavior that can be detrimental to survival (2). Thus, physiological mechanisms regulating alcohol intake are most likely highly conserved but remain sparsely described in humans.

Interestingly, acute alcohol exposure has little or no effect on plasma concentrations of hormones known to regulate energy homeostasis (e.g., insulin, C-peptide, glucagon, or the gut-derived glucoregulatory and/or satiety-promoting incretin hormones) (3), and various inflammation markers (4). In contrast, the liver-derived hormone fibroblast growth factor 21 (FGF21) has been shown to 1) be robustly secreted after alcohol intake in humans (3, 5) and 2) reduce ad libitum consumption of alcohol in animal studies (6, 7). Currently FGF21 is undergoing Phase 2 clinical trials for metabolic dysfunction-associated and alcohol-related liver disease (8). Whether other proteins are responsive to acute alcohol exposure remains uncertain.

Here, we used proteomics (9) to delineate the effect of acute alcohol exposure in healthy men on the plasma proteome, i.e., 1,462 different plasma proteins, with the aim of identifying proteins/biomarkers that are secreted in response to acute alcohol intake in humans. We hypothesized that FGF21 constitutes a major alcohol-responsive hormone, implicating it as a physiological regulator of alcohol consumption.

## Methods

### Participants

Plasma samples collected by Lanng et al. (3) from healthy men (n = 12) were examined. Key inclusion criteria were informed consent and body mass index (BMI) between 19 and 25 kg/m^2^. Key exclusion criteria were diabetes, anemia, liver diseases, nephropathy, and daily use of prescriptive medicine.

### Experimental procedures

This study was a post hoc analysis of a previously published study (3), which was approved by The Scientific-Ethical Committee of the Capital Region, Denmark (identification no. H-16026085) and registered at the Danish Data Protection Agency and Clinicaltrials.gov (Identifier: NCT03348371). The study was conducted according to the latest version of the Helsinki Declaration. The experimental procedures were previously described by Lanng et al. (3). In short, the participants met at the department after overnight fasting and received 0.7 g ethanol per kg body weight (20% alcohol solution (v/w)) as a bolus over 10 minutes through a nasogastric tube. The participants continued fasting until the end of the experimental day after four hours. Blood was collected in lithium heparin tubes (kept on ice) at baseline, and after 120 minutes, all tubes were centrifuged for 15 minutes at 2,000 *g* and 4°C. Samples were stored at −80°C until batch analysis.

### Analyses

The proteomic analysis was performed by Olink Proteomics, Uppsala Science Park, SE-751 83 Uppsala, Sweden. Olink uses a proximity extension assay based on oligonucleotide-linked antibody pairs with an affinity for each other. Oligonucleotide-linked antibody pairs were added to the plasma samples and when the antibodies are brought in proximity, the two unique oligonucleotides linked to the antibodies will be extended by a DNA polymerase and amplified exponentially. Quantitative real-time polymerase chain reaction (qPCR) amplifies and quantifies the oligonucleotides in the sample and serves as a surrogate marker of specific antigens that the antibodies recognize (10). The number of qPCR cycles is related to the protein concentration in the sample, and the relative concentration of each protein can be detected (9).

### Statistics

Quality control procedures were applied, correcting for batch effects, sample integrity, and technical performance. With 12 participants, we were powered (β=0.8) to detect a 65% increase in proteins whose levels in plasma vary similarly to FGF21. Generally, a 100% increase is used in proteomics studies to sort informative from non-informative hits.

## Results

As previously reported (3), we included 12 men with the following characteristics (mean ± SD): age 25 ± 3.9 years; body weight 77 ± 7.9 kg; BMI 23 ± 2.6 kg/m^2^; fasting plasma glucose 5.2 ± 0.39 mmol/l; HbA1c 5.3 ± 0.4%; plasma alanine aminotransferase 28 ± 11 U/l; plasma creatinine 87 ± 12 μmol/l. None of the participants were taking any medications regularly. All 12 participants completed the experimental day.

As published by Lanng et al., all participants started with a plasma ethanol concentration of 0 g/l. Plasma concentrations increased rapidly after alcohol administration and reached a peak concentration of 1.7 ± 0.3 g/l after 38 ± 18 minutes.

Out of 1,462 proteins analyzed, 11 significantly changed 120 minutes after alcohol intake. Seven proteins’ plasma concentrations decreased, and four proteins’ plasma concentrations increased after alcohol intake (Table 1 and Figure 1). The plasma concentration of FGF21 increased the most (10.5-fold) and most significantly (*P* < 0.000001) after alcohol intake. The second-greatest increase in plasma concentration after alcohol intake was insulin-like growth factor-binding protein 1 (*IGFBP1*) (2.1-fold, *P* = 0.000027), and the plasma concentration of semaphoring 4C increased third-most (*SEMA4C*) (1.4-fold, *P* = 0.000001) (Table 1 and Figure 1).

**Table 1.** Proteins exhibiting greatest differences in plasma concentrations after alcohol intake in men. Gene name, *P* value for the difference before vs. after alcohol intake, absolute change and fold increase/decrease after alcohol intake, and protein name are shown.

| Protein name (gene) | Difference | *P* value | Fold difference |
| --- | --- | --- | --- |
| Fibroblast growth factor 21 (*FGF21*) | 3.40 ± 0.20 | <0.000001 | 10.5-fold |
| Insulin-like growth factor-binding protein 1 (*IGFBP1*) | 1.10 ± 0.16 | 0.000027 | 2.1-fold |
| Semaphorin 4C (*SEMA4C*) | 0.62 ± 0.07 | 0.000001 | 1.4-fold |
| Left-right determination factor 2 (*LEFTY2*) | 0.29 ± 0.04 | 0.000046 | 1.2-fold |
| Myoglobin (*MB*) | -0.31 ± 0.03 | 0.000002 | 0.8-fold |
| C-C motif chemokine ligand 2 (*CCL2*) | -0.64 ± 0.12 | 0.000173 | -0.6-fold |
| Pancreatic polypeptide (*PYY*) | -0.42 ± 0.08 | 0.000343 | -0.7-fold |
| C-C motif chemokine ligand 4 (*CCL4*) | -0.50 ± 0.09 | 0.000204 | -0.7-fold |
| Motilin (*MLN*) | -0.62 ± 0.11 | 0.000119 | -0.7-fold |
| X-C motif chemokine ligand 1 (*XCL1*) | -0.26 ± 0.04 | 0.000104 | -0.8-fold |
| Matrix metalloproteinase 3 (*MMP3*) | -0.28 ± 0.05 | 0.000284 | -0.8-fold |

**Figure 1.**
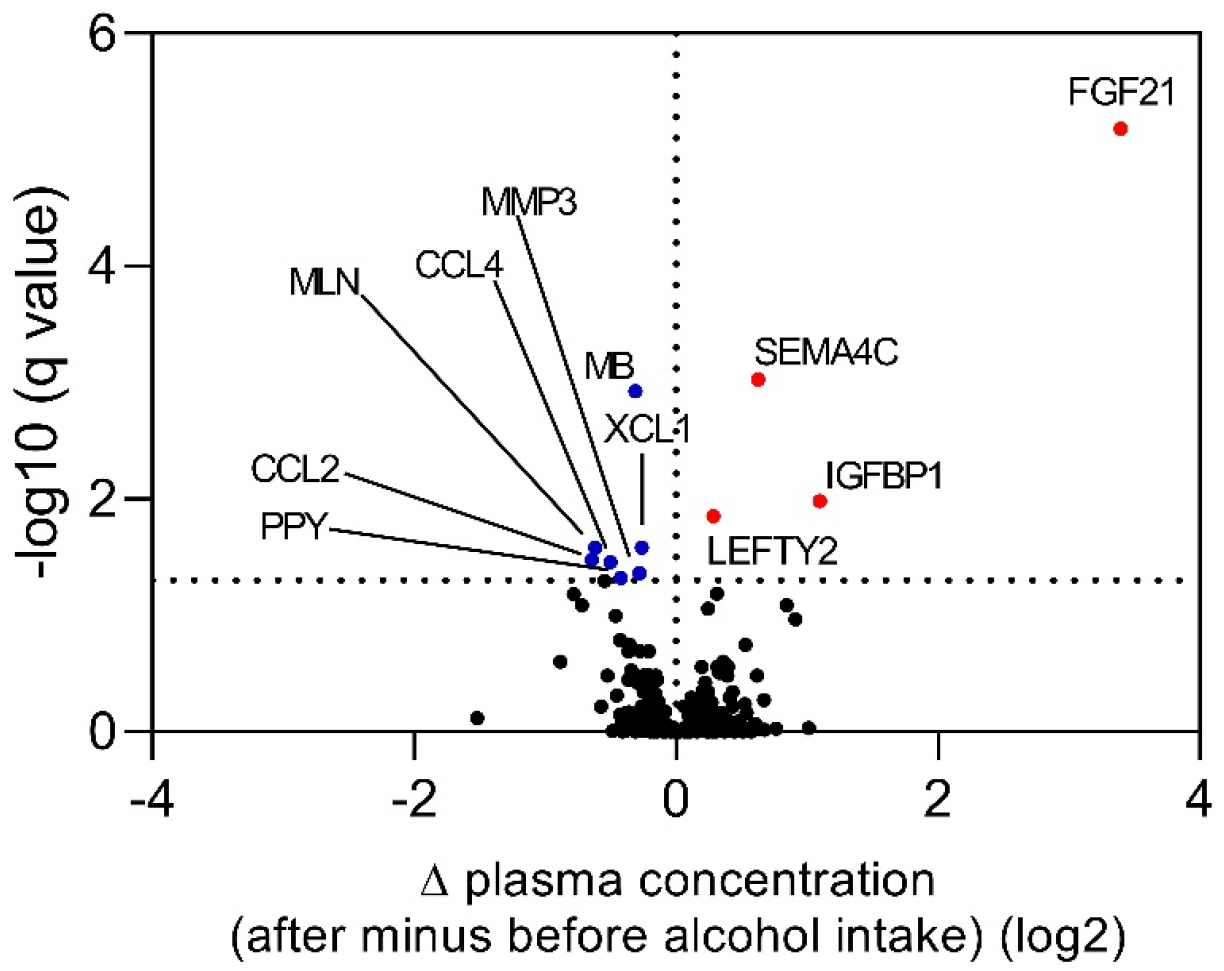
Volcano plot of changes in plasma protein concentrations from baseline to 120 minutes after alcohol intake and significant levels in healthy men. The horizontal dashed line marks statistical significance. The vertical line separates proteins that increased after alcohol intake (red dots) from those which decreased after alcohol intake (blue dots). Proteins exhibiting significant differences in plasma concentration after alcohol intake are labeled with their respective gene name.

## Discussion

Using plasma proteomic profiling, we show that acute alcohol exposure elicits a remarkably selective systemic protein response in healthy men. Despite profiling more than 1,400 proteins, the systemic proteomic response to acute alcohol exposure was remarkably limited, and only 11 proteins changed significantly two hours after alcohol administration, with FGF21 exhibiting by far the largest increase (>10-fold). These findings extend previous observations that alcohol stimulates FGF21 secretion (3, 5) by demonstrating that FGF21 is not simply one of many alcohol-responsive proteins but dominates the endocrine response to acute ethanol exposure in humans.

FGF21 is increasingly recognized as a liver-derived hormone that coordinates adaptive responses to nutritional and metabolic stress. Both experimental and clinical studies have shown robust increases in circulating FGF21 following acute alcohol intake (3, 11), while genetic studies have linked variants in the KLB gene, encoding the obligate co-receptor β-klotho, to alcohol consumption in humans (12, 13). Furthermore, pharmacological and genetic manipulation of the FGF21 pathway reduces alcohol preference in mice and non-human primates (6, 7), supporting the concept of a liver-brain endocrine feedback loop that limits excessive alcohol intake. Although our study was not designed to establish causality, the exceptional magnitude and specificity of the FGF21 response provide additional evidence that FGF21 represents a principal endocrine signal induced by alcohol exposure.

From this study an overall lack of widespread changes in the circulating proteome was seen. Despite exposure to an intoxicating dose of alcohol, fewer than 1% of the quantified proteins changed significantly. Moreover, the increase in FGF21 exceeded that of the second most responsive protein, IGFBP1, by approximately fivefold, emphasizing the unique nature of the FGF21 response. These findings are consistent with previous physiological studies demonstrating minimal acute effects of alcohol on classical metabolic hormones and inflammatory mediators (3, 4), suggesting that acute ethanol exposure triggers a highly specific rather than generalized endocrine response.

Our findings are also supported by recent targeted proteomic analyses. Stankevic and colleagues examined 92 inflammation-related proteins following binge alcohol exposure and likewise identified FGF21 as the most strongly induced circulating protein despite studying a substantially different protein panel and patient population (14). The concordance between these independent studies strengthens the evidence that FGF21 is a reproducible biomarker of acute alcohol exposure across diverse experimental settings.

Limitations of the study include the analysis was performed post hoc in a relatively small cohort of healthy young Caucasian men, and plasma proteins were measured at a single post-exposure time point. Consequently, the temporal dynamics of proteins with earlier or later responses may not have been captured, and the findings cannot necessarily be generalized to women, other ethnic groups, older individuals, or patients with alcohol use disorder. Validation in larger and more diverse cohorts will therefore be important.

In conclusion, we demonstrate that out of more than 1,400 proteins, only 11 changed significantly in plasma of 12 men two hours after alcohol exposure. FGF21 is the single-most responsive protein, showing a more than 10-fold increase in plasma concentration two hour after alcohol instillation, emphasizing FGF21’s potential role in regulating alcohol intake in humans.

## Conflict of interest statement

ARL and FKK are currently employed by Novo Nordisk, Bagsværd, Denmark; the present work was done independently from Novo Nordisk. MPG, SAJT, and LSG have no conflict of interest.

## Founding

This work was supported by Aase og Ejnar Danielsens Fond (grant number: 19-10-0069), A. P. Møller Fonden (grant number: 18-L-0126), Grosserer L. F. Foghts Fond (grant number: 21.519) and Savværksejer Jeppe Juhl og hustru Ovita Juhls mindelegat.

## Acknowledgments

We are thankful to all study subjects for their participation and lab-technicians Sisse M. Schmidt and Inass Al-Nachar (Center for Clinical Metabolic Research, Gentofte Hospital) for their invaluable help with experimental procedures.

## Data Availability Statement

Data is not available

## Notes

### Competing Interest Statement

Amelie Lanng and Filip Knop are employees of Novo. Work was conducted prior to employment.

## References

1. Carrigan MA, Uryasev O, Frye CB, Eckman BL, Myers CR, Hurley TD, et al. Hominids adapted to metabolize ethanol long before human-directed fermentation. Proc Natl Acad Sci U S A. 2015;112(2):458–63.

2. WHO. Global status report on alcohol and health 2028. WHO; 2018.

3. Lanng AR, Gasbjerg LS, Bergmann NC, Bergmann S, Helsted MM, Gillum MP, et al. Gluco-metabolic effects of oral and intravenous alcohol administration in men. Endocr Connect. 2019;8(10):1372–82.

4. Lanng AR, Gasbjerg LS, Bergmann NC, Gillum MP, Rehfeld JF, Helsted MM, et al. The effect of acute intragastric vs. intravenous alcohol administration on inflammation markers, blood lipids and gallbladder motility in healthy men. Alcohol. 2020;87:29–37.

5. Soberg S, Andersen ES, Dalsgaard NB, Jarlhelt I, Hansen NL, Hoffmann N, et al. FGF21, a liver hormone that inhibits alcohol intake in mice, increases in human circulation after acute alcohol ingestion and sustained binge drinking at Oktoberfest. Mol Metab. 2018;11:96–103.

6. Flippo KH, Trammell SAJ, Gillum MP, Aklan I, Perez MB, Yavuz Y, et al. FGF21 suppresses alcohol consumption through an amygdalo-striatal circuit. Cell Metab. 2022;34(2):317–28 e6.

7. Talukdar S, Owen BM, Song P, Hernandez G, Zhang Y, Zhou Y, et al. FGF21 Regulates Sweet and Alcohol Preference. Cell Metab. 2016;23(2):344–9.

8. Noureddin M, Frias JP, Neff GW, Lucas J, Behling C, Bedossa P, et al. Safety and efficacy of once-weekly efruxifermin versus placebo in metabolic dysfunction-associated steatohepatitis (HARMONY): 96-week results from a multicentre, randomised, double-blind,placebo-controlled, phase 2b trial. Lancet. 2025;406:719–30.

9. Cui M, Cheng C, Zhang L. High-throughput proteomics: a methodological mini-review. Lab Invest. 2022;102(11):1170–81.

10. Assarsson E, Lundberg M, Holmquist G, Bjorkesten J, Thorsen SB, Ekman D, et al. Homogenous 96-plex PEA immunoassay exhibiting high sensitivity, specificity, and excellent scalability. PLoS One. 2014;9(4):e95192.

11. Desai BN, Singhal G, Watanabe M, Stevanovic D, Lundasen T, Fisher FM, et al. Fibroblast growth factor 21 (FGF21) is robustly induced by ethanol and has a protective role in ethanol associated liver injury. Mol Metab. 2017;6(11):1395–406.

12. Schumann G, Liu C, O’Reilly P, Gao H, Song P, Xu B, et al. KLB is associated with alcohol drinking, and its gene product beta-Klotho is necessary for FGF21 regulation of alcohol preference. Proc Natl Acad Sci U S A. 2016;113(50):14372–7.

13. Sanchez-Roige S, Palmer AA, Fontanillas P, Elson SL, and Me Research Team tSUDWGotPGC, Adams MJ, et al. Genome-Wide Association Study Meta-Analysis of the Alcohol Use Disorders Identification Test (AUDIT) in Two Population-Based Cohorts. Am J Psychiatry. 2019;176(2):107–18.

14. Stankevic E, Israelsen M, Juel HB, Madsen AL, Angquist L, Aldiss PSJ, et al. Binge drinking episode causes acute, specific alterations in systemic and hepatic inflammation-related markers. Liver Int. 2023;43(12):2680–91.

